# Sex chromosomes control vertical transmission of feminizing *Wolbachia* symbionts in an isopod

**DOI:** 10.1101/747444

**Authors:** Thomas Becking, Mohamed Amine Chebbi, Isabelle Giraud, Bouziane Moumen, Tiffany Laverré, Yves Caubet, Jean Peccoud, Clément Gilbert, Richard Cordaux

**Author notes:** Corresponding author: Dr. Richard Cordaux. CG and RC are equal senior authors.

## Abstract

Microbial endosymbiosis is widespread in animals, with major ecological and evolutionary implications. Successful symbiosis relies on efficient vertical transmission through host generations. However, when symbionts negatively affect host fitness, hosts are expected to evolve suppression of symbiont effects or transmission. Here we show that sex chromosomes control vertical transmission of feminizing *Wolbachia* endosymbionts in the isopod *Armadillidium nasatum*. Theory predicts that the invasion of an XY/XX species by cytoplasmic sex ratio distorters is unlikely because it leads to fixation of the unusual (and often lethal or infertile) YY genotype. We demonstrate that *A. nasatum* X and Y sex chromosomes are genetically highly similar and YY individuals are viable and fertile, thereby enabling *Wolbachia* spread in this XY-XX species. Nevertheless, we show that *Wolbachia* cannot drive fixation of YY individuals because infected YY females do not transmit *Wolbachia* to their offspring, unlike XX and XY females. The genetic basis fits the model of a Y-linked recessive allele (associated with an X-linked dominant allele), in which the homozygous state suppresses *Wolbachia* transmission. Moreover, production of all-male progenies by infected YY females restores a balanced sex ratio at the host population level. This suggests that blocking of *Wolbachia* transmission by YY females may have evolved to suppress feminization, thereby offering a whole new perspective on the evolutionary interplay between microbial symbionts and host sex chromosomes.

## Introduction

Microbial endosymbiosis is widespread in animals, with major effects on host ecology and evolution [1, 2]. Successful symbiosis relies on efficient symbiont transmission through host generations. Vertical transmission occurs when symbionts are transferred from parent to offspring, usually through the maternal germ line [3, 4]. From a symbiont perspective, faithful maternal inheritance can be achieved by conferring benefits to hosts, such as nutritional provisioning [5], defense against natural enemies [6] and pathogen resistance [7, 8]. This transmission strategy leads to convergence of symbiont and host fitness. Conversely, symbionts may follow a selfish strategy consisting of favouring their transmission at the expense of host fitness. This is achieved through manipulation of host reproduction, which can result in highly distorted sex ratios [9, 10]. As balanced sex ratios are optimal for most nuclear genes owing to biparental inheritance, hosts are predicted to evolve suppression mechanisms to control symbiont effects or transmission [10, 11]. Genetic conflicts between sex ratio distorters, such as feminizing symbionts, and the rest of the genome are increasingly recognized as drivers of the evolution of host sex determination systems [10–16]. Here we take the reverse perspective and investigate whether host sex chromosome systems can influence symbiont transmission and dynamics.

In many animals, sex is determined by a locus located on sex chromosomes that is heterozygous in one sex (the heterogametic sex) and homozygous in the other sex (the homogametic sex). Two major types of sex chromosomes exist: male heterogamety (XY males and XX females) and female heterogamety (ZZ males and ZW females) [13, 15]. Sex chromosomes evolve from autosomes that acquired a sex-determining locus characterized by sex-specific inheritance. Subsequently, genomic regions around sex-determining loci often stop recombining, leading to gradual accumulation of nucleotide and structural variation and repetitive DNA [13,15,17–19]. This so-called degeneration process also causes the formation of pseudogenes and gene loss, resulting in increasing differentiation of sex chromosomes over time. This is well illustrated by the human X and Y sex chromosomes, which dramatically differ in size and gene content [19].

Arthropods host a diverse array of intracellular symbionts, including bacteria (such as *Wolbachia*, *Cardinium*, *Rickettsia*, *Spiroplasma* and others) and unicellular eukaryotes (microsporidia), that are able to distort host sex ratios towards females [9, 10]. The evolutionary impact of cytoplasmic sex ratio distorters on host sex determination systems has been particularly investigated in terrestrial isopods (crustaceans). This is because both female and male heterogametic systems of sex chromosomes are found in this speciose taxonomic group [20] and many species are naturally infected with *Wolbachia* bacteria [21, 22]. *Wolbachia* are intracellular, maternally inherited endosymbionts of arthropods, often acting as reproductive parasites that manipulate host reproduction to favour infected females in host populations [23]. In terrestrial isopods, *Wolbachia* is best known as a sex ratio distorter due to its ability to feminize genetic males into phenotypic females [10,24–26]. For example, the presence of feminizing *Wolbachia* in *Armadillidium vulgare* leads to highly female-biased progenies [27, 28] because symbionts override the ZZ/ZW system of sex chromosomes [29, 30]. As a result, genetic ZZ male embryos develop as phenotypic ZZ females when infected by *Wolbachia*. Theoretical models predict the extinction of the W sex chromosome in *A. vulgare* lines infected by *Wolbachia* [31, 32] and empirical evidence verified this prediction [33, 34]. Thus, all individuals of *Wolbachia*-infected lineages end up with ZZ sex chromosomes at equilibrium, those inheriting *Wolbachia* develop as females and those lacking *Wolbachia* develop as males. This phenomenon is known as cytoplasmic sex determination [10,24–26]. In sum, invasion of a ZZ/ZW species by feminizing *Wolbachia* is not problematic because it leads to the fixation of the ZZ genotype, which is the natural male genotype.

In sharp contrast with ZZ/ZW heterogamety, the conditions allowing feminizing *Wolbachia* to spread in an XY/XX heterogametic system are much more stringent. Indeed, theoretical work predicts that infection of an XY/XX system by a cytoplasmic sex ratio distorter should drive the loss of the X chromosome and concomitantly lead to fixation of the YY genotype at equilibrium [31]. In this case, YY individuals inheriting *Wolbachia* should develop as females and those lacking *Wolbachia* should develop as males. However, YY is not a standard genotype and it may often be lethal or infertile, owing to Y chromosome degeneration causing the loss of essential genes [13,15,17–19]. Therefore, XY/XX heterogamety is expected to be incompatible with invasion of feminizing *Wolbachia* symbionts, unless the X and Y chromosomes are not substantially divergent, i.e. they both carry all vital loci and are genetically very similar. To the best of our knowledge, this prediction has never been explored empirically.

Here, we investigated the interplay between sex chromosomes and *Wolbachia* symbionts in *Armadillidium nasatum*, a terrestrial isopod species related to *A. vulgare* [20]. Analogous to *A. vulgare*, a feminizing *Wolbachia* strain (*w*Nas) is naturally present in *A. nasatum* [21,35,36]. However, unlike *A. vulgare*, chromosomal sex determination follows XY/XX heterogamety in *A. nasatum* [20, 35]. This result was established using an original strategy consisting of experimentally reversing young genetic females into phenotypic males, crossing them with their sisters and analyzing sex ratios of the resulting progenies [20, 35]. Here, based on whole-genome sequencing, pedigree analyses and simulations, we show that *Wolbachia* endosymbionts can spread in *A. nasatum* populations because *A. nasatum* sex chromosomes are genetically highly similar and YY males and females are both viable and fertile. Nevertheless, *Wolbachia* cannot drive the loss of the X chromosome because infected YY females do not transmit *Wolbachia* to their offspring, unlike XX and XY females. As infected YY females produce all-male progenies, a balanced sex ratio is maintained at the host population level despite the presence of feminizing *Wolbachia*, suggesting that blocking of *Wolbachia* transmission by YY females may have evolved to suppress feminization.

## Results

### De novo assembly and annotation of male (XY) A. nasatum genome

We sequenced the male (XY) genome of an *A. nasatum* line derived from wild animals sampled in Thuré (France) in 2008. Males and females from this line have been consistently producing progenies with balanced sex ratios in our laboratory. In addition, sex reversal and crossing experiments demonstrated that sex determination in this line follows male heterogamety [20]. Absence of *Wolbachia* endosymbionts in the individuals selected for sequencing was confirmed by PCR.

We assembled the *A. nasatum* genome using a hybrid approach combining short paired-end Illumina reads and long PacBio reads (S1 Table). The initial assembly was processed through polishing, decontamination and scaffolding by a transcriptome assembly. The final assembly had a total length of 1,223,175,971 bp. It was composed of 25,196 contigs and scaffolds (hereafter collectively referred to as “contigs”) with an N_50_ of 86,284 bp, and containing barely no (0.01%) undetermined nucleotides (Table 1). Genome completeness assessment using Benchmarking Universal Single Copy Orthologs (BUSCO 3.0.1) [37] revealed that 1,001 of 1,066 (94%) conserved specific arthropod genes were present in the assembly (Table 1). Furthermore, transcriptome assembly alignment on the genome assembly yielded 99.0% of transcripts longer than 1 kb aligned. Thus, we have obtained a reliable assembly of the XY male genome from *A. nasatum*.

**Table 1.** Summary information of *Armadillidium nasatum* genome assembly.

| Genome assembly features | Assembly figures |
| --- | --- |
| Assembly statistics |  |
| Number of contigs and scaffolds | 25,196 |
| Total size | 1,223,175,971bp |
| Longest contig/scaffold | 877,941bp |
| Number of contigs and scaffolds >1 kb | 25,166 |
| N<sub>50</sub> contig size | 75,116bp |
| N<sub>50</sub> scaffold size | 86,284bp |
| Undetermined nucleotides | 0.01% |
| G+C content | 25.77% |
| Analysis of BUSCO genes |  |
| Complete genes | 974/1066 (91.4%) |
| Complete and single-copy genes | 932/1066 (87.4%) |
| Complete and duplicated genes | 42/1066 (3.9%) |
| Fragmented genes | 27/1066 (2.5%) |
| Missing genes | 65/1066 (6.1%) |

The annotation of the *A. nasatum* assembly included 14,636 predicted gene models with an average length of 9,422 bp and representing 11.3% of the total assembly length (S2 Table). Among these predicted genes, 9,281 (63.4%) had Blastp hits to the UniProt-SwissProt database (release September 2016) and 9,459 (64.6%) had InterProScan hits to Pfam domains (version 30.0) and 6,137 (41.9%) had Gene Ontology terms. Repeats accounted for 64.6% (or 790 Mb) of the *A. nasatum* assembly (S3 Table). Specifically, transposable elements and simple tandem repeats accounted for 40.6% (or 496 Mb) and 23.1% (or 283 Mb) of the genome. The annotated genome sequence of *A. nasatum* is available in DDBJ/ENA/GenBank under accession number SEYY00000000. The version described in this article is version SEYY01000000.

### High genetic similarity of the X and Y chromosomes

To investigate the extent of genomic differentiation between sex chromosomes, we searched the *A. nasatum* assembly for contigs containing Y-specific sequences. We mapped short paired-end Illumina reads generated for XY males and XX females (S1 Table) onto the *A. nasatum* assembly and performed a Chromosome Quotient analysis [38]. This analysis consists of comparing the ratios of female-to-male sequencing depths (CQ) for each contig, with the expectations that: (i) Y-specific contigs should be mapped by male reads only (CQ ∼0), (ii) X-specific contigs should be mapped by female reads at twice the sequencing depth of male reads (CQ ∼2), and (iii) autosomal contigs should be mapped at similar sequencing depths by female and male reads (CQ ∼1). The resulting frequency distribution of CQ scores was unimodal and centered at CQ ∼1 (mean: 1.03, median: 1.04), with no peak at CQ scores of ∼0 and ∼2 (Fig 1a). In addition, we used the mapping-free Y chromosome Genome Scan (YGS) method [39], which computes the proportion of single copy k-mers for each contig of the genome assembly of the heterogametic (male) sex that are unmatched to sequencing reads of the homogametic (female) sex. Y-specific contigs are expected to be unmatched by female reads (YGS ∼100%) while autosomal and X-linked contigs are expected to be matched entirely (YGS ∼0%). The resulting frequency distribution of YGS scores indicated that most contigs have very low YGS scores (mean: 14.5%, median: 11.2%) and very few contigs had high YGS scores (i.e. only 43 contigs with YGS≥80% and just 2 with YGS≥90%) (Fig 1b). Thus, the CQ and YGS analyses consistently indicated that the *A. nasatum* assembly mostly contains autosomal contigs and very few contigs containing X- and Y-specific sequences. Intersecting the results of the CQ and YGS analyses identified only 78 out of the 25,196 contigs of the assembly as containing putative Y-specific sequences, despite the use of permissive thresholds, i.e. CQ≤0.35 and YGS≥35% (Fig 1c). The 78 contigs comprised a total length of 1,327 kb, corresponding to just 0.1% of the *A. nasatum* genome assembly (S4 Table). While most contigs lacked any gene (65/78, or 83%), a total of 20 genes (out of 14,636 genes in the assembly) were annotated from 13 contigs (S4 Table). They constitute candidate master genes for sex determination in *A. nasatum*. Thus, the Y-specific region of the *A. nasatum* genome is extremely small (around one megabase) and contains very few genes, leading to the conclusion that the X and Y chromosomes are poorly differentiated at the genomic and genic levels.

**Fig 1.**
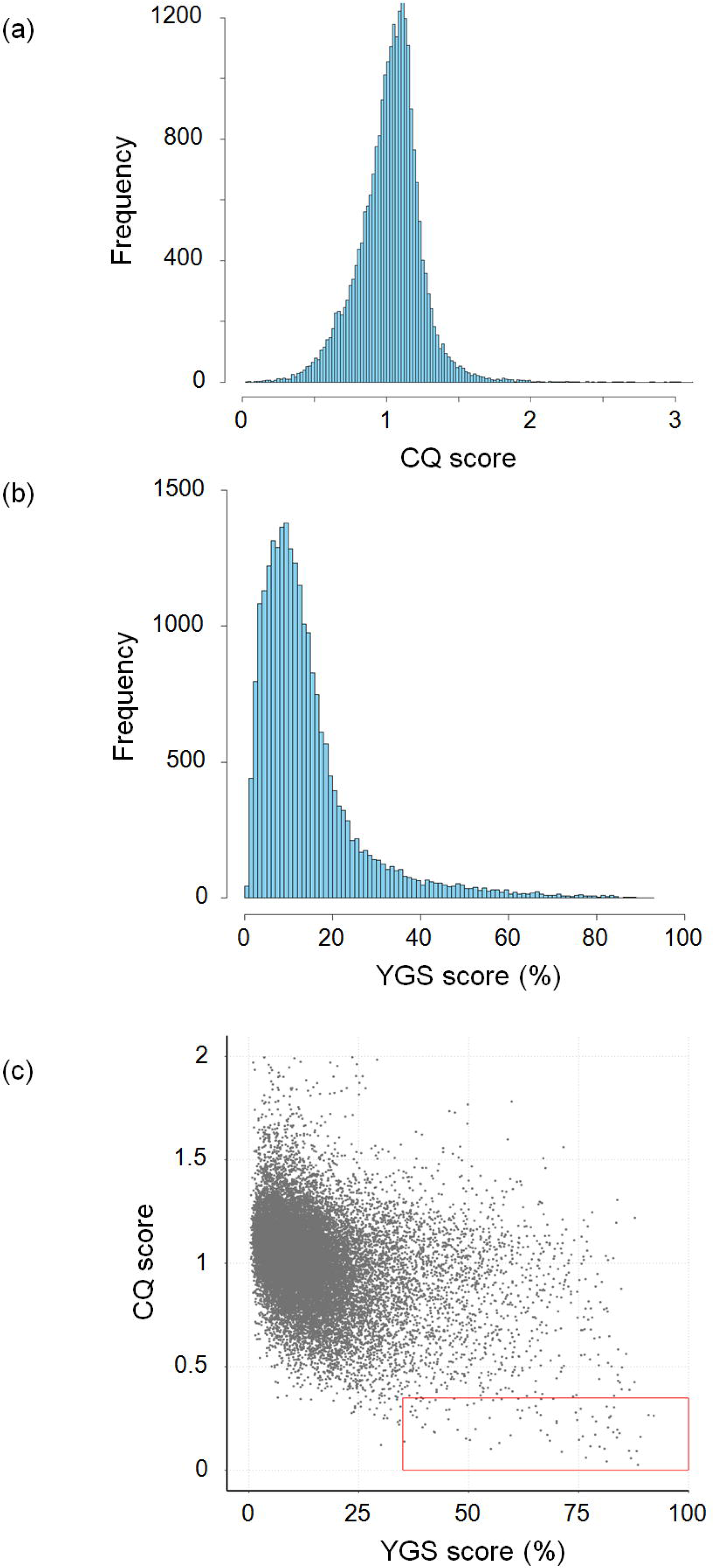
Identification of sex-specific contigs in the *Armadillidium nasatum* genome assembly. Frequency distribution of CQ (a) and YGS (b) scores calculated for each contig and scaffold of the assembly. (c) Comparison of CQ and YGS scores. Each dot corresponds to one contig or scaffold (those with CQ>2 are not represented). The red box contains 78 contigs with CQ≤0.35 and YGS≥35%.

To investigate sex chromosome differentiation at the molecular level, we analyzed the patterns of sequence divergence and repeat content in the 78 contigs containing putative Y-specific sequences relative to the other contigs of the *A. nasatum* genome. First, we analyzed the density of single nucleotide polymorphisms (SNP) in the 78 contigs (after removal of hemizygous regions, to focus on regions with orthologs on the X chromosome) versus the other contigs of the assembly. We calculated a median SNP density of 2.61 SNP/kb between allelic X/Y regions of the 78 contigs, compared to 2.55 SNP/kb across all other contigs of the assembly (S1 Fig). Thus, we observed a slight but non-significant (Mann-Whitney two-sided U test, U=969,520, p=0.88) excess of SNP density in allelic X/Y regions relative to other regions of the genome. In addition, the repeat content in the 78 contigs was slightly higher than (median 69.6%), but not significantly different (Mann-Whitney bilateral U test, U=1,048,600, p=0.28) from the repeat content of all other contigs (median 67.8%) (S1 Fig). Furthermore, even when focusing on 25 of the 78 contigs that were independently validated as Y-linked by PCR (see below), there was no significant difference with all other contigs of the assembly, both in terms of SNP density (median 3.91 SNP/kb, Mann-Whitney two-sided U test, U=371,560, p=0.11) and repeat content (median 65.3%, Mann-Whitney two-sided U test, U=301,380, p=0.73) (S1 Fig). Thus, the potential presence of (autosomal) false positives among the 78 contigs containing putative Y-specific sequences has not affected our results. In sum, our results indicated that *A. nasatum* sex chromosomes present patterns of molecular evolution that are quite similar to those of other genomic regions, with a slight elevation of SNP density and repeat content in contigs containing putative Y-specific sequences that is consistent with expectations for sex-specific sequences of the genome [40]. We conclude that the X and Y chromosomes of *A. nasatum* are highly similar not only at the genomic and genic levels, but also at the molecular level.

### YY individuals are viable and fertile

To test for the existence and, if so, viability and fertility of YY individuals, we tracked Y chromosome inheritance in a *Wolbachia*-infected *A. nasatum* pedigree. First, we established robust Y-specific molecular markers using the putative Y-specific contigs identified previously. Reliable PCR assays were successfully designed for 42 of the 78 contigs containing putative Y-specific sequences, most of which (25/42) exhibited male-specific amplification with DNA samples from males and females closely related to those used for genome sequencing (S4 Table). However, only 9 of the 25 confirmed markers displayed male-specific amplification when tested in more distantly related individuals from the same population and in individuals from two other populations. Lack of sex specificity of a subset of markers in some populations indicated that recombination has occurred between some of the tested contigs and the sex-determining locus. This result provided further evidence that the Y-specific region of the genome is extremely small in *A. nasatum*. Concomitantly, the universal male specificity of 9 markers across all tested populations demonstrated that these *A. nasatum* populations possess a homologous Y chromosome. Hence, we used a subset of these robust markers to track the Y chromosome in an *A. nasatum* pedigree spanning three generations.

The pedigree was initiated by crossing a male with a female naturally infected by the feminizing *Wolbachia* strain *w*Nas, which produced an F1 progeny comprising 17 sisters (Fig 2). Two F1 sisters were then crossed with genetic males (XY), to produce two F2 progenies: I-F2-1 comprising 24 females and I-F2-2 comprising 32 males. *Wolbachia* infection was tested in the 62 individuals of the pedigree (except the F0 father), indicating that all females carried *Wolbachia* while all males lacked *Wolbachia* infection, as expected. Next, we used previously designed Y-specific markers to assess the presence of the Y chromosome in the 62 individuals (except the F0 father). In the I-F2-1 progeny, half of the females showed amplification of the Y markers (indicating they are XY or YY) and half did not (indicating they are XX). The F2-1 mother carried XX chromosomes, as she did not amplify the Y markers. As all I-F2-1 offspring necessarily carry one X chromosome from their mother, the 11 I-F2-1 females amplifying the Y markers must be XY. In the I-F2-2 progeny, all individuals amplified the Y markers (indicating they are XY or YY), as did their mother. Given that the F2-2 father is XY, if the F2-2 mother was XY, we would have expected 25% XX individuals in the I-F2-2 progeny, but there was none. Thus, the I-F2-2 mother must be YY, which is the only genotype that can explain that the entire I-F2-2 progeny carries at least one Y chromosome. Given the parental genotypes, we predicted the I-F2-2 progeny is composed of half XY males and half YY males. Finally, the two F1 sisters being XX and YY, it followed that the F0 father and mother are XY. To independently assess these predictions, we devised a quantitative PCR assay measuring Y chromosome dose relative to autosomes (S2 Fig). This assay confirmed the predicted genotypes of the F0 and F1 individuals we tested and the composition of the I-F2-2 progeny (15 XY and 17 YY individuals) (Fig 2, S5 Table). In sum, the resolution of sex chromosome genotypes in the pedigree demonstrated that the YY genotype is viable both as male and as female in *A. nasatum*, as is the XY genotype. Moreover, the YY and XY genotypes are both fertile as females. Finally, we show below that the YY genotype is also fertile as male.

**Fig 2.**
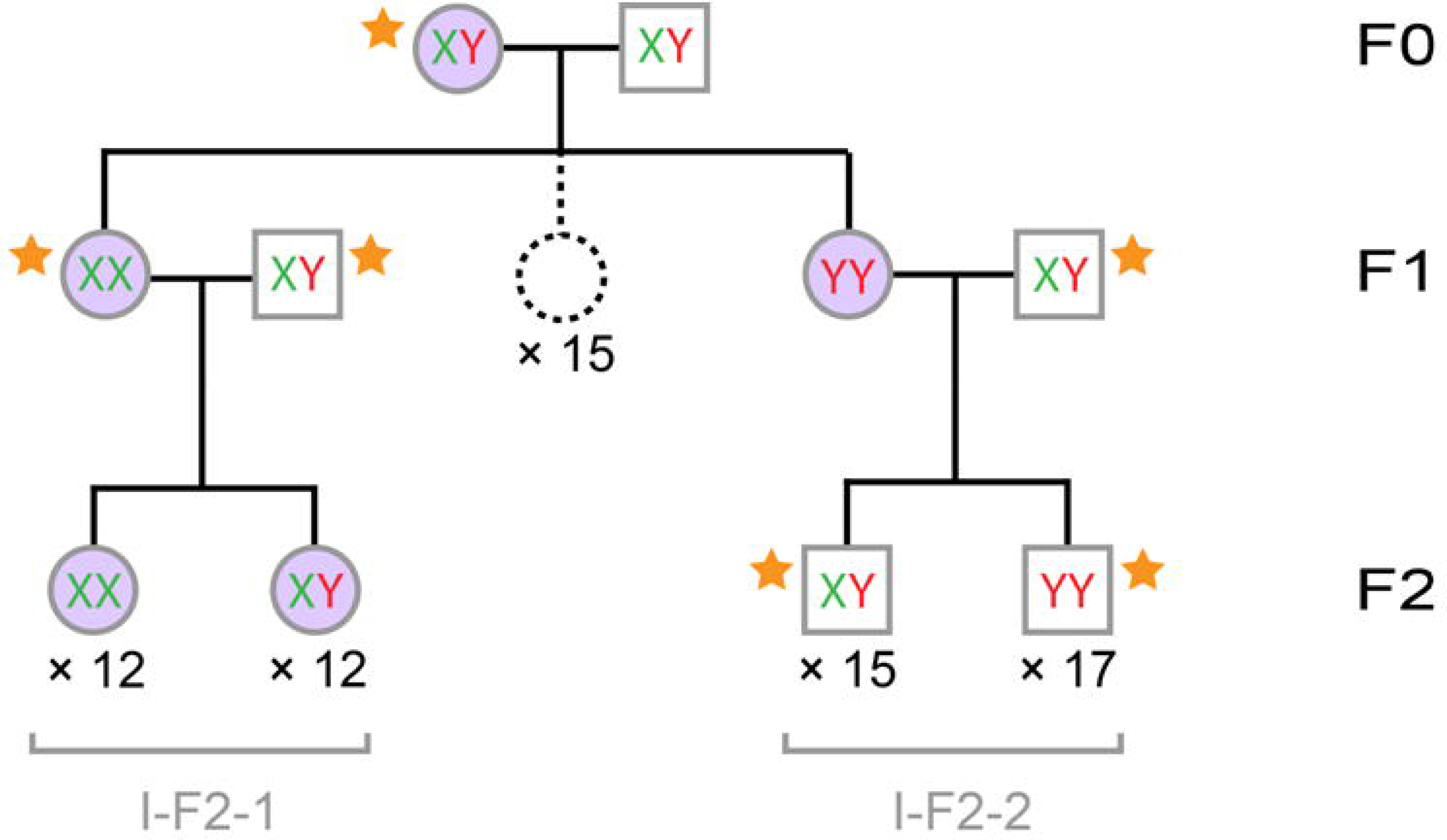
*Armadillidium nasatum* pedigree used to track inheritance of the Y chromosome and *Wolbachia*. The pedigree spans three generations (F0-F2) and is comprised of 62 individuals (35 males and 27 females) for which sex chromosome genotype (XX, XY or YY) was identified and 15 F1 females not included in the molecular analyses (dotted circle). Males are shown as squares and females as circles. Individuals carrying *Wolbachia* are shown in purple. Progeny IDs are shown in grey. Sex chromosome genotype of individuals marked with an orange star (34 males and 2 females) was also assessed with a quantitative PCR assay.

### Blocking of Wolbachia transmission by YY females prevents X chromosome loss

Surprisingly, the *Wolbachia*-infected YY mother in the previous pedigree analysis produced an all-male progeny (I-F2-2) entirely lacking *Wolbachia* endosymbionts (Fig 2). This is unexpected because infection by feminizing *Wolbachia* endosymbionts is usually associated with highly female-biased progenies, due to the maternal transmission of *Wolbachia* to usually >80% of the offspring [10,24–26]. To test whether lack of *Wolbachia* transmission from mother to I-F2-2 progeny was a random event or was due to the unusual YY maternal genotype, we extended our previous pedigree analysis to span five generations (S3 Fig) and analysed a second, independent pedigree spanning four generations (S4 Fig). In total, 20 families (defined as father, mother and progeny) were included in the two pedigrees, representing 799 individuals (252 males and 547 females). We tested 464 individuals for the presence of *Wolbachia* (Table 2). As expected, most tested females were infected by *Wolbachia* (180/214) while no tested male was (0/250). Based on the patterns of Y chromosome amplification and following a reasoning similar to that described in the previous section, we were able to infer sex chromosome genotypes for most analysed individuals. In addition, we independently verified sex chromosome genotypes of the parents of the 20 families using the aforementioned quantitative PCR assay (S5 Table). The sex chromosome genotypes of the 20 mothers included 11 XX, 5 XY and 4 YY females (S3 and S4 Fig, Table 2). All tested fathers were XY males, except the father of the II-F2-1 progeny which was a YY male. Interestingly, this individual demonstrated that the YY genotype is viable and fertile as male.

**Table 2.** Composition and frequency of *Wolbachia* infection in 20 families of *Armadillidium nasatum*. *Wolbachia* frequency in progenies was calculated as the *Wolbachia* frequency observed among tested females weighted by female proportion in the progenies (as all males lack *Wolbachia*).

| Progeny ID | Parental genotypes |  | Progeny size | Female number | Female proportion | *Wolbachia* presence in males |  | *Wolbachia* presence in females |  | *Wolbachia* frequency |
| --- | --- | --- | --- | --- | --- | --- | --- | --- | --- | --- |
|  | Mother | Father |  |  |  | Nb. tested | Nb. infected | Nb. tested | Nb. infected |  |
| I-F1-1 | XY | XY | 17 | 17 | 100% | 0 | 0 | 3 | 3 | 100% |
| I-F2-1 | XX | XY | 24 | 24 | 100% | 0 | 0 | 24 | 24 | 100% |
| I-F2-2 | YY | XY | 32 | 0 | 0% | 32 | 0 | 0 | 0 | 0% |
| I-F2-3 | XX | XY | 52 | 52 | 100% | 0 | 0 | 2 | 2 | 100% |
| I-F3-1 | XY | XY | 68 | 68 | 100% | 0 | 0 | 12 | 12 | 100% |
| I-F3-2 | XX | XY | 33 | 33 | 100% | 0 | 0 | 10 | 8 | 80% |
| I-F3-3 | XX | XY | 59 | 59 | 100% | 0 | 0 | 12 | 12 | 100% |
| I-F3-4 | XX | XY | 40 | 40 | 100% | 0 | 0 | 10 | 9 | 90% |
| I-F4-1 | XX | XY | 50 | 39 | 78% | 11 | 0 | 15 | 12 | 62% |
| I-F4-2 | XX | XY | 41 | 24 | 59% | 17 | 0 | 13 | 0 | 0% |
| I-F4-3 | XX | XY | 32 | 10 | 31% | 22 | 0 | 10 | 2 | 6% |
| I-F4-4 | XX | XY | 16 | 16 | 100% | 0 | 0 | 10 | 9 | 90% |
| I-F4-5 | XX | XY | 51 | 51 | 100% | 0 | 0 | 15 | 15 | 100% |
| I-F4-6 | XY | XY | 42 | 19 | 45% | 23 | 0 | 13 | 7 | 24% |
| II-F1-1 | XX | Y? | 13 | 13 | 100% | 0 | 0 | 2 | 2 | 100% |
| II-F2-1 | XY | YY | 60 | 60 | 100% | 0 | 0 | 60 | 60 | 100% |
| II-F2-2 | XY | XY | 20 | 20 | 100% | 0 | 0 | 1 | 1 | 100% |
| II-F3-1 | YY | XY | 28 | 0 | 0% | 28 | 0 | 0 | 0 | 0% |
| II-F3-2 | YY | XY | 42 | 0 | 0% | 42 | 0 | 0 | 0 | 0% |
| II-F3-3 | YY | XY | 57 | 0 | 0% | 57 | 0 | 0 | 0 | 0% |

*Wolbachia* transmission rate from mother to offspring differed significantly between XX, XY and YY mothers, considering all 20 families (Kruskal-Wallis test, χ²=8.84, p=0.01) or only the 16 families in which *Wolbachia* presence was tested in ≥10 individuals (Kruskal-Wallis test, χ²=7.91, p=0.02). *Wolbachia* transmission rate was high (generally ≥80%) and did not differ significantly between XX and XY mothers (Dunn’s post-hoc tests, p=0.73 for 20 families and p=0.56 for 16 families) (Fig 3). By contrast, *Wolbachia* infected none of the 159 offspring of the 4 YY mothers. This result cannot be ascribed to *Wolbachia* transmission to offspring and subsequent selective death of embryos carrying *Wolbachia* infection because progeny size did not differ significantly between XX, XY and YY mothers (Kruskal-Wallis tests, χ²=0.18, p=0.91 for 20 families and χ²=3.32, p=0.19 for 16 families) (Table 2). Instead, the YY genotype is associated with a lack of *Wolbachia* transmission from mother to offspring. As a result, *Wolbachia* transmission rate differed significantly between YY mothers and both XX mothers (Dunn’s post-hoc tests: p=0.006 for 20 families and p=0.01 for 16 families) and XY mothers (Dunn’s post-hoc tests: p=0.008 for 20 families and p=0.01 for 16 families) (Fig 3). Consequently, as all offspring inherited a Y chromosome from their YY mother but no *Wolbachia* endosymbiont, they all developed as males (Table 2). Importantly, YY mothers originated from the two independent pedigrees we analysed (S3 and S4 Fig), indicating that the observed pattern is robust to *A. nasatum* genetic background.

**Fig 3.**
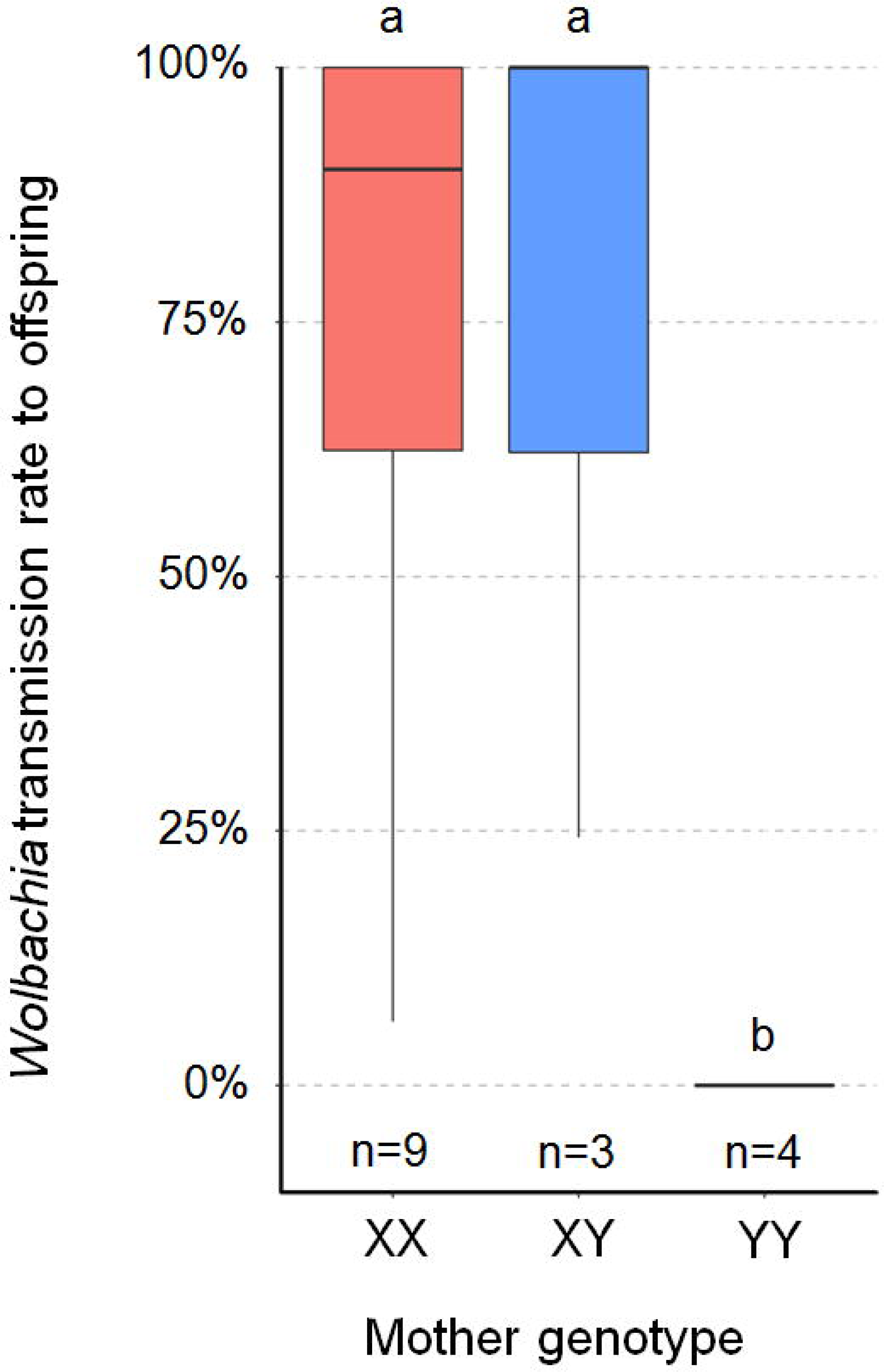
Boxplot of *Wolbachia* transmission rate from mother to offspring (measured as the frequency of *Wolbachia*-carrying individuals in each progeny) according to mother’s sex chromosome genotype. The analysis is based on 16 progenies (n) for which *Wolbachia* presence was tested in ≥10 individuals. Thick lines and boxes depict median and interquartile range, respectively. Whiskers are bounded to the most extreme data point within 1.5 the interquartile range. Plots marked with the same letter (a, b) are not statistically different from each other (Kruskal-Wallis test, followed by pairwise comparison Dunn test).

As YY females exclusively produce males, only XX and XY females can produce female offspring in *Wolbachia*-infected lines of *A. nasatum*. Thus, the X chromosome is necessarily transmitted to at least a subset of each progeny of these *Wolbachia*-infected females. To evaluate consequences on sex chromosome frequencies in the long term, we extended Taylor’s theoretical work [31] to enable the transmission rate of a cytoplasmic sex ratio distorter to vary depending on female sex chromosome genotype. Specifically, we simulated the equilibrium frequencies of X and Y chromosomes in a population of a diploid genetic model containing a distorter with transmission rates of α, α and 0 in XX, XY and YY females (with α varying from 0 to 1), respectively, to reflect our empirical results. Under these conditions, equilibrium frequencies are 25% and 75% for the X and Y chromosomes for α<0.55 (Fig 4a). In such cases, the distorter is lost from the population at equilibrium and the population is comprised of XY males and XX females in equal proportions. By contrast, for higher α values, equilibrium frequencies of the X and Y chromosomes vary in opposite directions, to the extent that the Y chromosome becomes more frequent than the X chromosome, but X equilibrium frequency is always ≥12.5%, whatever α (Fig 4a). Thus, a consequence of the lack of transmission of the distorter by YY females is that the X chromosome cannot be lost from the population. Another consequence evidenced by our simulations is that the equilibrium frequency of the distorter never exceeds 50%, whatever α (Fig 4a). Remarkably, a balanced sex ratio is maintained in the population at equilibrium, despite infected females individually producing progenies highly biased towards either females (XX and XY mothers) or males (YY mothers) (Fig 4b). This is in sharp contrast with the highly female-biased sex ratios at equilibrium expected for α>0.5 when the distorter is transmitted by all three types of females [31] (Fig 4b).

**Fig 4.**
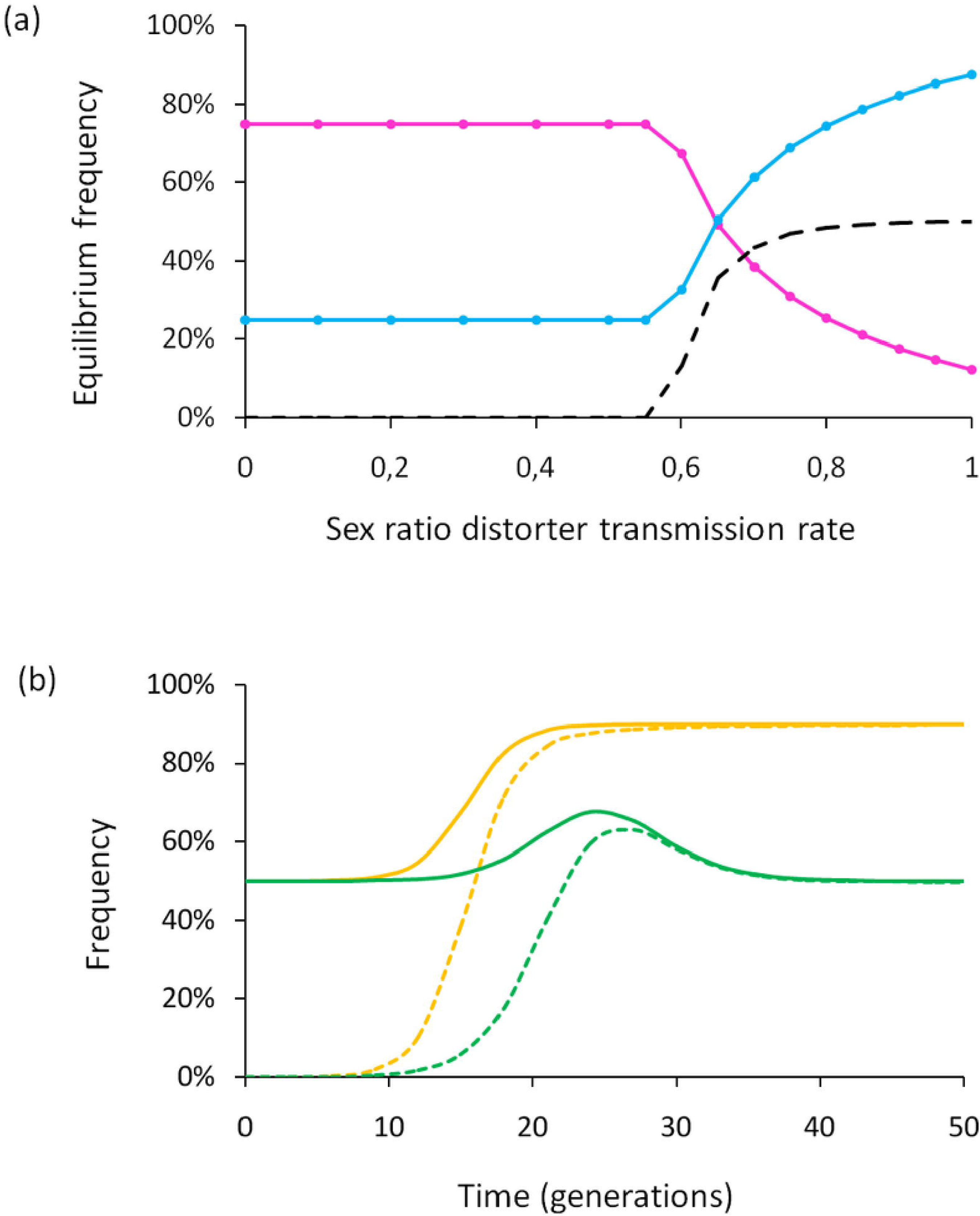
Evolutionary consequences of a cytoplasmic sex ratio distorter at the population level. (a) Equilibrium frequencies of X chromosome (pink line), Y chromosome (blue line) and distorter (dashed line), according to distorter transmission rate in XX and XY females (YY females do not transmit the distorter). (b) Evolution of the frequencies of females (solid lines) and individuals carrying the distorter (dashed lines) through time, with a distorter transmission rate of 0.9. Green: YY females do not transmit the distorter (only XX and XY females do); orange: all females transmit the distorter.

## Discussion

We sequenced the genome of the terrestrial isopod *A. nasatum*. With an N_50_ of 86 kb, high completeness (BUSCO=94%) and near absence of unidentified nucleotides (0.01%), our ∼1.2 Gb assembly is among the best assembled of all large-genome crustaceans sequenced to date [30]. Our results establish that *A. nasatum* X and Y chromosomes are genetically highly similar. The Y-specific region of the genome may be as small as one megabase and it contains very few genes. Even at nucleotide resolution, sequence divergence and repeat content indicate very limited sex chromosome differentiation. This result explains why the unusual YY genotype is viable in this species. The underlying causes of the very high similarity of *A. nasatum* sex chromosomes include young evolutionary age, ongoing recombination or both. In any event, this high similarity was likely instrumental to the establishment of feminizing *Wolbachia* infection in the male heterogametic isopod *A. nasatum*.

An important outcome of the infection of *A. nasatum* by feminizing *Wolbachia* is the unusual production of YY individuals and XY females. This situation challenges the classical model of sex chromosome evolution by affecting chromosome effective population size (N_e_) and recombination patterns. Classically, recombination arrest occurs between the X and Y chromosomes, leading to a drastic reduction of Y chromosome N_e_ to one third of X chromosome N_e_ and one fourth of autosome N_e_ (because there is one Y chromosome dose for three X chromosome doses and four autosome doses per mating pair). Lowered N_e_ results in enhanced genetic drift and concomitant reduction of selection efficiency, leading the Y chromosome to accumulate deleterious mutations and degenerate [13,15,17–19]. However, the existence of YY individuals and XY females implies that the standard expectation of sex chromosome N_e_ does not apply to *A. nasatum* in the presence of *Wolbachia*. Indeed, our simulations indicate that X chromosome N_e_ should stabilize at about one fourth of Y chromosome N_e_ and about one fifth of autosome N_e_ at observed *Wolbachia* transmission rates (Fig 4a). Thus, X chromosome N_e_, not Y chromosome N_e_, is predicted to be dramatically reduced in *A. nasatum* genetic backgrounds infected by feminizing *Wolbachia*. Furthermore, the occurrence of YY individuals opens the possibility of Y-Y recombination and more efficient selection, which may prevent the accumulation of deleterious mutations on the Y chromosome. By contrast, XX individuals are rare according to our simulations (<2%), thereby reducing the opportunity of X-X recombination. This suggests that the X chromosome, rather than the Y chromosome, may accumulate deleterious mutations and degenerate in the long term. Another possibility to consider is that recombination patterns may differ between sexes (heterochiasmy) in some species, so that the X and Y chromosomes may not recombine simply because males do not recombine (whereas females do) [15]. If so, the existence of XY females in *A. nasatum* as a consequence of *Wolbachia* infection would open the possibility of X-Y recombination in phenotypic females, as previously reported in frogs [41]. Unfortunately, male and female patterns of recombination are unknown in terrestrial isopods. Their investigation therefore represents a perspective for future research. In any event, there appears to be ample scope for microbial symbionts to drive the molecular evolution of their host sex chromosomes.

Remarkably, infected YY females did not transmit *Wolbachia* to any offspring, as confirmed in two independent pedigrees. An evolutionary consequence is that the YY genotype cannot become fixed (nor the X chromosome lost) in infected *A. nasatum* lines. Lack of *Wolbachia* transmission by YY females also raises the question of the underlying mechanism controlling vertical transmission of the symbionts. In sharp contrast with the YY genotype, the alternative XX and XY genotypes were associated with similarly high rates of *Wolbachia* transmission in both pedigrees. Thus, the genetic basis of *Wolbachia* transmission control fits the model of a recessive allele linked to the Y chromosome (associated with an X-linked dominant allele), in which the homozygous state blocks *Wolbachia* transmission. It has been shown previously that within-host microbial interactions can enhance or reduce symbiont vertical transmission [42]. This applies to *Wolbachia*, which vertical transmission is prevented by *Asaia* bacteria in *Anopheles* mosquitoes [43]. Host nuclear genotype is also an important factor that can affect symbiont vertical inheritance. For example, *Wolbachia* maternal transmission is strongly reduced by actin in *Drosophila* [44] and the *wds* gene in *Nasonia* [45]. Our results show that sex chromosome genotype represents yet another case of host nuclear control over symbiont transmission. *Wolbachia* symbionts were previously found to prevent sex chromosome transmission in *Eurema mandarina* butterflies [46]. Here we show that sex chromosomes can prevent *Wolbachia* transmission in *A. nasatum*, thereby offering a whole new perspective on the molecular interplay between feminizing symbionts and host sex chromosomes.

The mutation causing the suppression of *Wolbachia* transmission in *A. nasatum* may have existed in the standing pool of sex chromosome variants prior to *Wolbachia* infection, having evolved either neutrally or under natural selection. Alternatively, the mutation may have evolved as an adaptation subsequent to *Wolbachia* infection. An excellent empirical example supporting the latter hypothesis is offered by the butterfly *Hypolimnas bolina*, in which suppression of *Wolbachia*-mediated male killing was found to have evolved in Southeast Asia secondarily to *Wolbachia* spread in the Indo-Pacific region [47]. While infection by feminizing symbionts normally leads to highly female-biased sex ratios, the ability of infected YY females to impede *Wolbachia* transmission in *A. nasatum* results in the production of all-male progenies, due to the systematic inheritance of a maternal Y chromosome. Interestingly, this situation restores balanced sex ratios at the population level, as indicated by our simulations (Fig 4) and consistent with empirical evidence [35, 48]. This opens the possibility that blocking of *Wolbachia* transmission by YY females may have evolved to suppress feminization and ensuing sex ratio biases, as predicted by sex ratio selection [10–12,15].

Indeed, strong sex ratio biases towards females imposed by feminizing symbionts induce nucleo-cytoplasmic conflicts with most nuclear genes, which are biparentally inherited and optimally benefit from balanced sex ratios. In principle, conflict resolution may be achieved by restoring balanced sex ratios at the parental level, through selection of females producing balanced sex ratios [49]. Alternatively, resolution may occur at the population level, through selection of females producing male-biased sex ratios to compensate for the female-biased progenies of infected females [49]. YY females in *A. nasatum* may represent an original example of conflict resolution at the population level. Several cases of nuclear suppression preventing the action or transmission of sex ratio distorting symbionts have been reported [11], including in the isopod *A. vulgare* [50, 51]. A distinguishing feature of symbiont suppression in *A. nasatum* is its connection with sex chromosomes.

## Materials & Methods

### Genome sequencing and assembly

All *A. nasatum* individuals used for sequencing were from our inbred laboratory line ANa, which is originally derived from wild animals caught in Thuré, France, in 2008. Specifically, we used XY genetic males and XX genetic females descended from a single pair of grandparents of the family ANa2 (according to the crossing scheme shown in S5 Fig) to minimize heterozygosity. Total genomic DNA was extracted using the QiagenDNeasy Blood and Tissue Kit, according to the protocol for animal tissues (3 h of incubation in proteinase K at 56°C and 30 min of RNase treatment at 37°C). Absence of *Wolbachia* endosymbionts in all samples was confirmed by PCR using the *ftsZ* and *wsp* markers [21, 36]. Short paired-end libraries with ∼200 bp insert sizes were sequenced with the Illumina HiSeq2000 technology (S1 Table). In addition, PacBio RS II sequencing (P6C4 chemistry) was performed to obtain long sequencing reads (S1 Table). Accession numbers for Illumina and PacBio sequence datasets are provided in S1 Table.

Sequencing reads were subjected to quality control and filtering as described previously [30]. Male Illumina and PacBio sequencing reads were used in a hybrid strategy to assemble the male (XY) genome of *A. nasatum*, as described previously [30]. A summary of the workflow we used (including assembly, polishing, contaminant removal and scaffolding) is shown in S6 Fig. Genome assembly completeness was evaluated using benchmarking for universal single copy orthologs (BUSCO, version 3.0.1) [37], with the arthropod profile library (-l arthropoda) composed of 1,066 arthropod core genes. Repeat identification and gene annotation were performed as described previously [30].

### Identification and analyses of contigs containing putative Y-specific sequences

Contigs containing putative Y-specific sequences in the *A. nasatum* assembly were identified using the Chromosome Quotient (CQ) [38] and Y chromosome Genome Scan (YGS) [39], as described previously [30]. The maximum CQ score was set to 0.35 to retain contigs as Y-specific candidates, as recommended by CQ authors [38]. The minimum YGS score was set to 35% to retain contigs as Y-specific candidates. This threshold was selected to account for the high repetitive nature of the *A. nasatum* genome. To identify heterozygous SNPs in the contigs of the male genome assembly, we applied the Genome Analysis ToolKit (GATK) pipeline (version 3.8-0-ge9d806836) [52], as described previously [30].

PCR assays were designed on the candidate contigs as follows (S4 Table). Primers were designed with Primer-BLAST [53] in unique regions of the contigs and primer specificity was checked using Blastn (version 2.2.30+) [54] by aligning primers to the unmasked *A. nasatum* assembly. PCR reactions were carried out in 25 μL with 5 μL of buffer 5×, 0.5 μL of dNTPs (2.15 mM), 0.7 μL of each primer (10 μM), 0.25 μL of Taq polymerase 5 u/μL, and 1 μL of DNA. PCRs were conducted using the following temperature cycle: 3 min at 94°C for initial denaturation, followed by 35 cycles of 30 s at 94°C, 30 s at 48/50/52°C (depending on primer annealing temperature) and 1 min at 72°C. The final elongation step was 10 min at 72°C. PCR tests were then conducted in three successive steps: (i) test on one male and three pools of two females of the ANa2 family (used for genome sequencing); (ii) test on six males and twelve females from other families of our ANa laboratory line, and (iii) test on two males and a pool of three females from Beauvoir-sur-Niort population (France) and on two males and a pool of three females from Piriápolis population (Uruguay). After each step, loci amplifying in all males and no female were retained for the next step. PCR tests targeting the autosomal *18S rRNA* [30] and mitochondrial *COI* [55] genes were used as positive controls in all samples. Absence of *Wolbachia* endosymbionts was also confirmed in all samples by PCR using the *ftsZ* and *wsp* markers [21, 36].

### Pedigree construction and analyses

We generated two independent *A. nasatum* pedigrees spanning five (pedigree I, S3 Fig) and four (pedigree II, S4 Fig) generations (*A. nasatum* has a generation time of one year). We started in 2013 with two F0 female founders isolated from a laboratory cage population (NASw) initiated in 2008 from wild animals (caught in Thuré, France) infected by the feminizing *Wolbachia* strain *w*Nas [36]. Each F0 female produced an F1 progeny (resulting from mating with unknown F0 males from the cage population). Each year, two to six females were selected from the previous generation and crossed with XY genetic males from our ANa laboratory line (except the father of the II-F2-1 progeny, which was selected from the NASw cage population and could carry XY or YY sex chromosomes). Pedigrees I and II were composed of 14 and 6 families, respectively.

At each generation, total genomic DNA was extracted from the progenitors (except the F0 male founders and individuals which died prematurely) and from males and females of their progenies. Presence or absence of *Wolbachia* was tested by PCR using the *ftsZ* and *wsp* markers [21, 36]. We also used two of the previously designed Y-specific markers (contig12740 and contig18908) to assess the presence of the Y chromosome in the individuals, interpreting PCR results as follows: amplification indicating the individual is XY or YY and lack of amplification indicating that the individual is XX. PCR tests targeting the autosomal *18S rRNA* [30] and mitochondrial *COI* [55] genes were used as positive controls in all samples. Based on PCR amplification patterns and pedigree structures, we were able to infer sex chromosome genotypes (XX, XY or YY) for most analyzed individuals.

To independently assess sex chromosome genotypes (XX, XY or YY), we developed a quantitative PCR assay measuring Y chromosome abundance relative to autosomes. Y-specific primers were designed within the Y-specific PCR amplicon previously validated in contig10349 (primer sequences: 5’-CCCTACACAGCATACTTGACAG and 5’-CAGGTGCTCCTTCAGAGAAAC, product size: 129 bp). Contig10349 abundance was normalized against *EF2* gene abundance [56]. *EF2* is a single-copy gene in the *A. nasatum* assembly located in autosomal contig8976 (YGS=8.8% and CQ=1.09). PCR reactions were run in duplicates in a 480 Light-Cycler (Roche) using SYBR Green I assays, under the following conditions: 10 min at 95°C and 45 cycles of [10 sec at 95°C, 10 sec at 60°C, 20 sec at 72°C]. A melting curve (65°C to 97°C) was recorded at the end of each reaction to check PCR product uniqueness. Reaction mixture consisted of 0.5 μL of each primer (10 μM), 5μL of Fast SYBR-Green Master Mix (Roche) 3 μL of bi-distilled water and 1 µL of extracted DNA. Fluorescence crossing points (C_t_) were calculated with the second derivative maximum method using the LightCycler 1.5 software. PCR amplification efficiency was determined with a calibration curve for each primer pair. Only PCR reactions producing a single product and with C_t_≤35 cycles were considered. Relative abundance of the Y-specific marker relative to autosomal marker was calculated as 2^-ΔCt^, where ΔC_t_ = C_t(Y chromosome)_ –C_t(autosome)_. XX, XY and YY genotypes corresponded to 2^-ΔCt^ values of ∼0, ∼0.5 and ∼1, respectively.

### Simulation of sex chromosome frequencies

To evaluate the evolutionary impact of cytoplasmic sex ratio distorters on sex chromosome frequencies, we extended Taylor’s theoretical work [31] to enable the transmission rate of the distorter to vary according to female sex chromosome genotypes. Specifically, we simulated the frequencies of the X chromosome, the Y chromosome, the distorter and females in a population of a diploid genetic model with a distorter whose transmission rate can vary according to the sex chromosome genotype of females (i.e. XX, XY and YY). We used the following equations:

Genotypes

- Males: G_1_=XY; G_3_=YY; F_M_=Males frequency
- Females: G_2_=XX; G_4_=XX.Wo+; G_6_=XY.Wo+; G_8_=YY.Wo+; F_F_=Females frequency

Gametes

- Males: P_1_=Y; P_3_=X
- Females: P_2_=X; P_4_=Y; P_6_=X.Wo+; P_8_=Y.Wo+

*Wolbachia* transmission rates by females

- t_X_ by XX.Wo+; t_XY_ by XY.Wo+; t_Y_ by YY.Wo+

Gametes production formula

- Male gametes: 

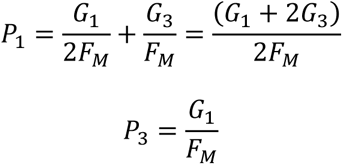

- Female gametes: 

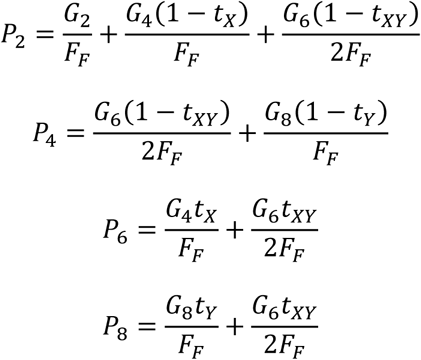

Next generation genotypes production formula

- Males: 

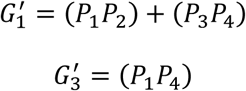

- Females: 

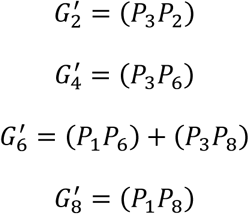
 with 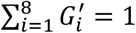 and 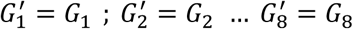 at equilibrium.

## Legends to supporting information

Fig. S1. Box plots of SNP density (a) and repeat proportion (b) in the 78 computationally-inferred, putative Y-linked contigs, a subset of 25 of the 78 contigs that were independently validated as Y-linked by PCR, and all the other contigs of the assembly. Thick lines and boxes depict median and interquartile range, respectively. Whiskers are bounded to the most extreme data point within 1.5 the interquartile range. Open circles represent outliers.

Fig. S2. Characterization of sex chromosome genotypes (XX, XY or YY) of *A. nasatum* individuals based on a quantitative PCR assay. Y chromosome to autosome ratios were calculated for 60 individuals and compared to expected ratios: 1 for YY individuals (corresponding to 18 males and 3 females), 0.5 for XY individuals (28 males and 5 females) and 0 for XX individuals (6 females). Thick lines and boxes depict median and interquartile range, respectively. Whiskers are bounded to the most extreme data point within 1.5 the interquartile range.

Fig. S3. *Armadillidium nasatum* pedigree I used to track inheritance of the Y chromosome and *Wolbachia*. The pedigree spans five generations (F0-F4) and is comprised of 572 individuals (119 males and 453 females), 269 of which were included in molecular analyses (individuals not included in the molecular analyses are shown in dotted circles). Males are shown as squares and females as circles. Individuals carrying *Wolbachia* are shown in purple. Progeny IDs are shown in grey. Sex chromosome genotype of individuals marked with an orange star was also assessed with a quantitative PCR assay.

Fig. S4. *Armadillidium nasatum* pedigree II used to track inheritance of the Y chromosome and *Wolbachia*. The pedigree spans four generations (F0-F3) and is comprised of 226 individuals (132 males and 94 females), 196 of which were included in molecular analyses (individuals not included in the molecular analyses are shown in dotted circles). Males are shown as squares and females as circles. Individuals carrying *Wolbachia* are shown in purple. Progeny IDs are shown in grey. Sex chromosome genotype of individuals marked with an orange star was also assessed with a quantitative PCR assay.

Fig. S5. Crossing scheme used to obtain *Armadillidium nasatum* individuals whose genomic DNA was extracted for Illumina (orange) and PacBio (red) sequencing. Males are shown as squares and females as circles.

Fig. S6. Workflow of the hybrid strategy used for assembling the *Armadillidium nasatum* genome.

Table S1. Characteristics of *Armadillidium nasatum* sequencing datasets generated in this study.

Table S2. Annotation statistics of the *Armadillidium nasatum* genome assembly.

Table S3. Repeat content of the *Armadillidium nasatum* genome.

Table S4. Characteristics of the 78 contigs of the *Armadillidium nasatum* assembly considered as Y-specific candidates.

Table S5. PCR results for Y chromosome and *Wolbachia* analyses of 72 *Armadillidium nasatum* individuals. nt: not tested because male individual originating from *Wolbachia*-free line, hence necessarily XY (except II-F1-1 father from *Wolbachia*-infected line, hence XY or YY). Y chromosome to autosome ratio was calculated as 2^-ΔCt^.

