## Supplementary figures and images for "Sex chromosomes control vertical transmission of feminizing *Wolbachia* symbionts in an isopod"

### Suppl Figure S1

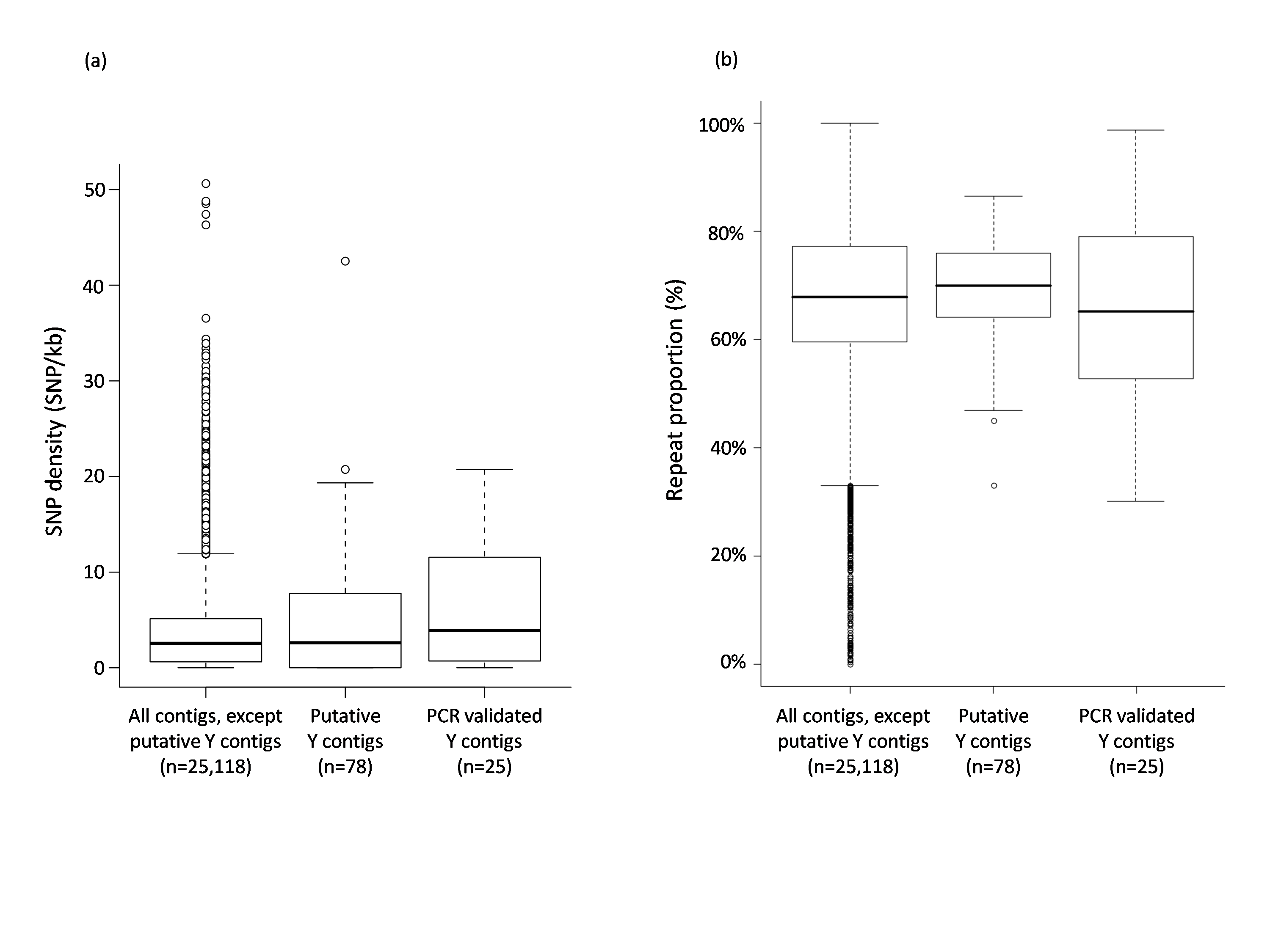

### Suppl Figure S2

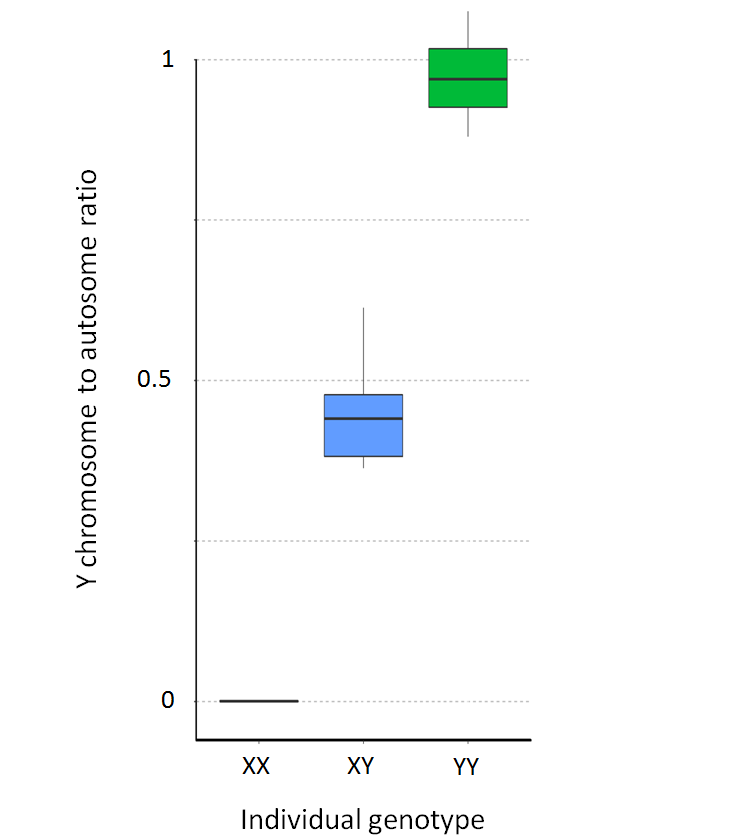

### Suppl Figure S3

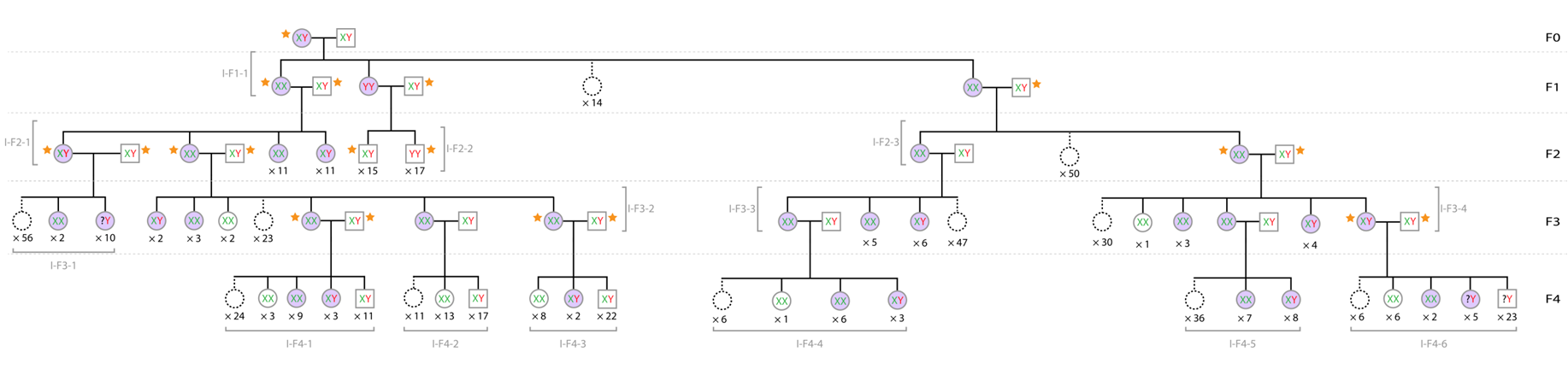

### Suppl Figure S4

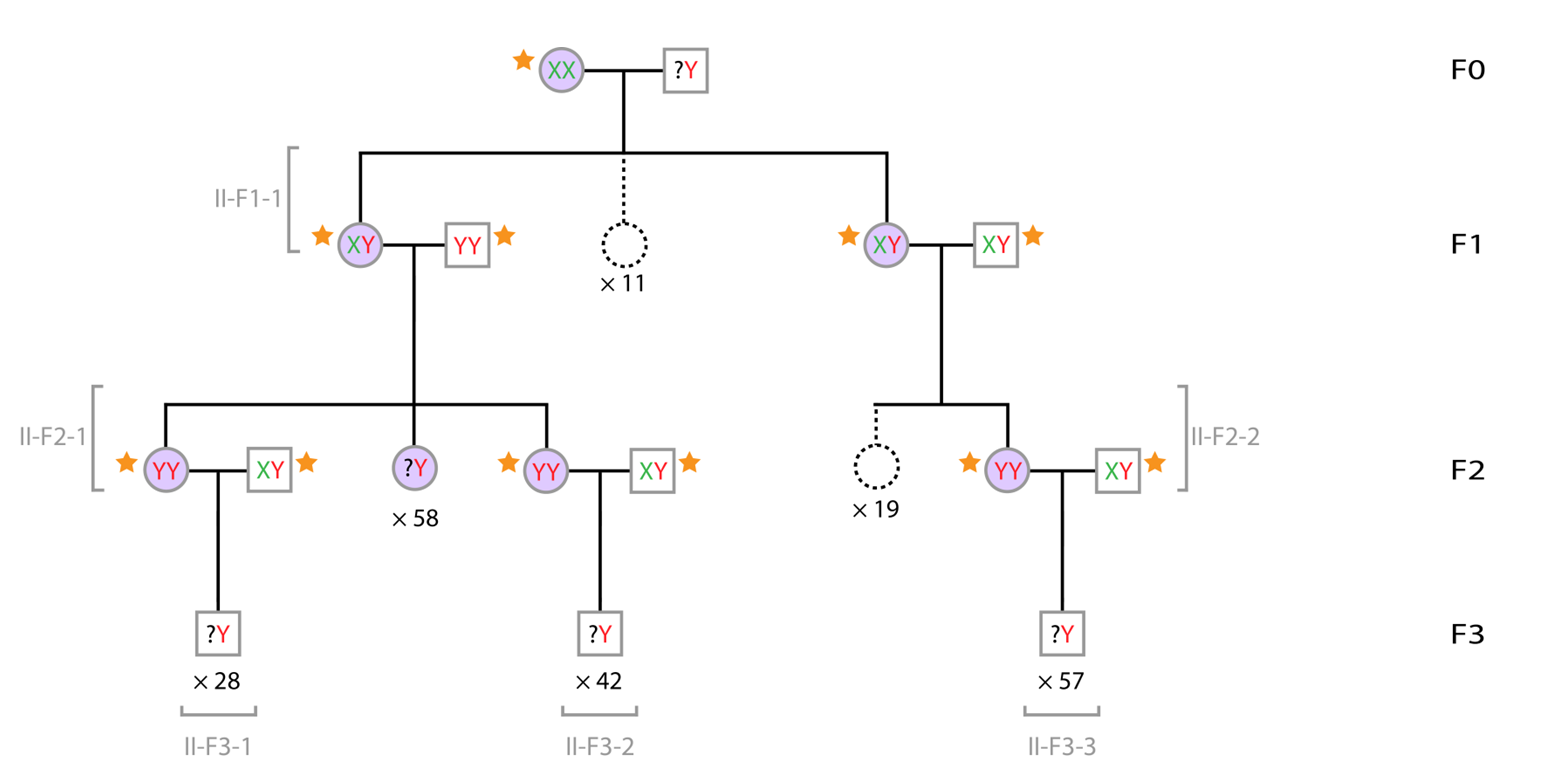

### Suppl Figure S5

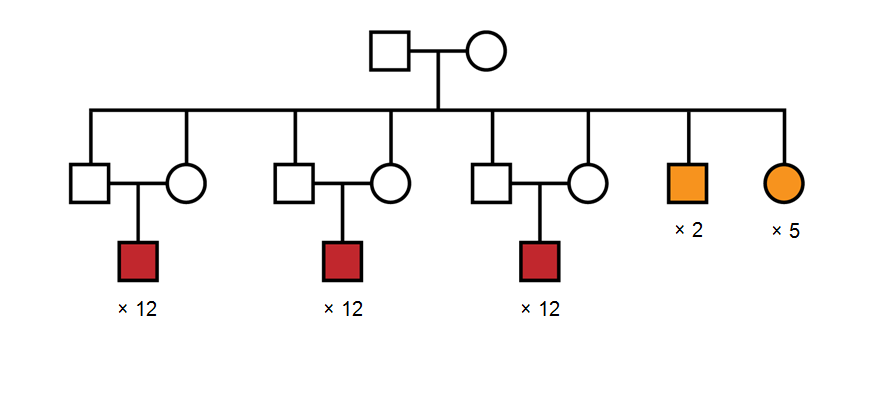

### Suppl Figure S6

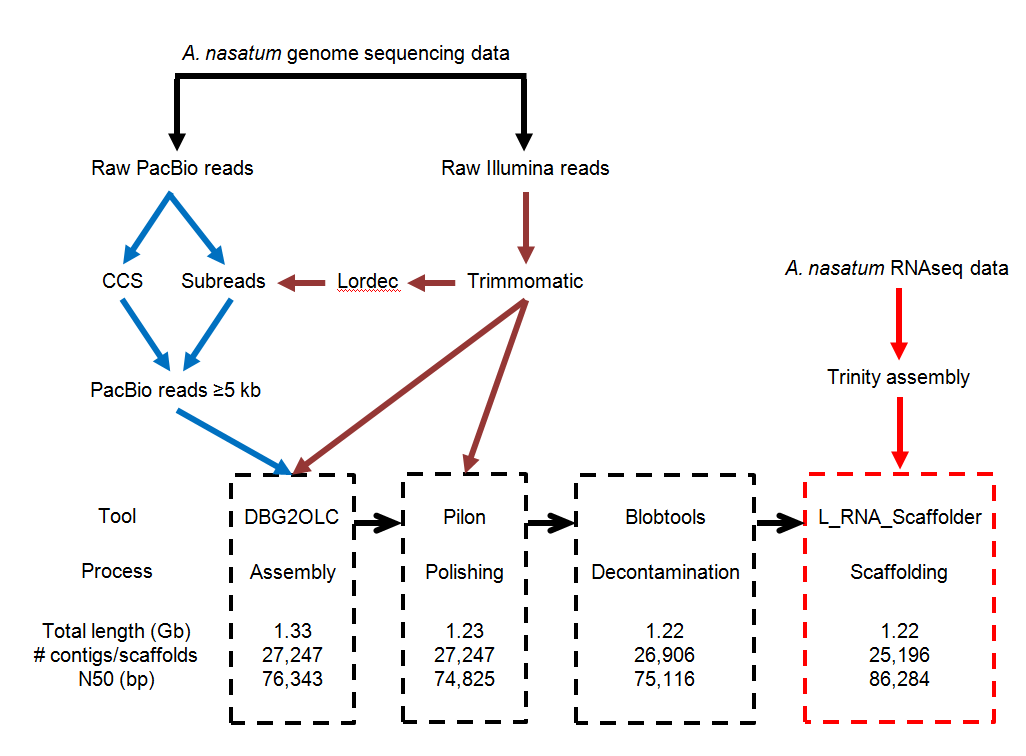
